# A Neural Network Based Algorithm for Dynamically Adjusting Activity Targets to Sustain Exercise Engagement Among People Using Activity Trackers

**DOI:** 10.1101/775908

**Authors:** Ramin Mohammadi, Amanda Jayne Centi, Mursal Atif, Stephen Agboola, Kamal Jethwani, Joseph C Kvedar, Sagar Kamarthi

**Affiliations:** Department of Mechanical and Industrial Engineering, Northeastern University, Boston, United States; Connected Health Innovation, Partners Healthcare, Boston, United States; Department of Dermatology, Massachusetts General Hospital, Boston, United States; Harvard Medical School, Harvard University, Boston, United States

**Keywords:** Activity trackers, exercise engagement, dynamic activity targets, neural network, activity target prediction, machine learning

## Abstract

It is well established that lack of physical activity is detrimental to overall health of an individual. Modern day activity trackers enable individuals to monitor their daily activity to meet and maintain targets and to promote activity encouraging behavior. However, the benefits of activity trackers are attenuated over time due to waning adherence. One of the key methods to improve adherence to goals is to motivate individuals to improve on their historic performance metrics. In this work we developed a machine learning model to dynamically adjust the activity target for the forthcoming week that can be realistically achieved by the activity-tracker users. This model prescribes activity target for the forthcoming week. We considered individual user-specific personal, social, and environmental factors, daily step count through the current week (7 days). In addition, we computed an entropy measure that characterizes the pattern of daily step count for the current week. Data for training the machine learning model was collected from 30 participants over a duration of 9 weeks. The model predicted target daily count with mean absolute error of 1545 steps. The proposed work can be used to set personalized goals in accordance with the individual’s level of activity and thereby improving adherence to fitness tracker.

## Introduction

Studies have reported the efficacy of physical activity in reducing the risk of disease. However, physical inactivity is on rise in the US population [1]. Considering that physical inactivity was the fourth leading cause of mortality in 2016 [2], there is much emphasis on developing effective methods to increase and maintain healthy levels of physical activity. One promising solution is wearable fitness trackers that enable individuals to monitor their activity levels and patterns to manage their health [3].

It is reported that about 20% of the general health-tracking population uses smart devices such as medical gadgets, mobile phone apps, or online tools to track their health data [4]. The use of technology to objectively monitor physical activity is associated with higher levels of activity [5]. However, the potential benefits derived from the use of physical activity trackers are challenged by the limited and transient adoption of these devices, which necessarily require sustained use to exert their intended effect. The continued engagement with fitness trackers is an issue that warrants further investigation [5].

Previous study [6] found two factors that are associated with the adoption and sustained use of physical activity trackers: (a) the number of digital devices owned by the participants and (b) the use of activity fitness trackers and other smart devices by if the participants’ family members bode well for the increased use of activity trackers. Studies have demonstrated that motivational factors are associated with physical activity levels [1]. Lack of time, fatigue and a dislike for exercise are some of the barriers that reflect a lack of motivation to engage in physical activity [1]. It is reported that adjustment to targets for activity levels are likely to enhance the users’ commitment to physical activity and engagement with fitness trackers [7].

With the rise of machine learning and availability of the activity tracker data, it is possible to create a model that can learn from users’ behavior and dynamically adjust the activity target for a forthcoming. Machine learning has been broadly applied within healthcare fields such as but not limited to cancer staging, risk assessment and drug recommendation systems [8]. Researchers studied the accuracy of activity trackers for energy expenditure assessment [9]. Having an automated personal trainer developed by data mining techniques can be useful for an amateur sportsman who cannot afford a personal trainer [10]. Similarly, effective feedback methods can be used for helping athletes and coaches [11].

Although, there have been some attempts at studying the benefit of activity trackers, there is a room for improvement. This study, to the best of our knowledge, is the first of its kind to develop a machine learning method to dynamically adjust the activity target for activity trackers. Machine learning techniques such as, but not limited to, lasso regression (LASSO), ridge regression (RIDGE), Bayesian ridge regression (BRIDGE), neural networks (NNs), random forest regressions (RF) and support vector regressions (SVR) have been used for prediction in the medical field [12].

Feature selection is an important step for improving the model performance [13]. Prior to applying machine learning techniques, it is essential to study the data to find features that might negatively or positively affect the model [8]. In this work we applied two feature selection techniques separately, principal component analysis (PCA) [14] and recursive feature elimination (RFE), with support vector machine (SVM) [15]. These are described in the methods section.

We compared predictive models developed over (a) all features, (b) features generated by PCA, (c) features selected by RFE and (d) features found by authors in the previous work. Figure 1 shows the flow diagram of the study highlighting the key steps. Later in the methods section, we explain in detail the stages for model development.

**Figure 1:**
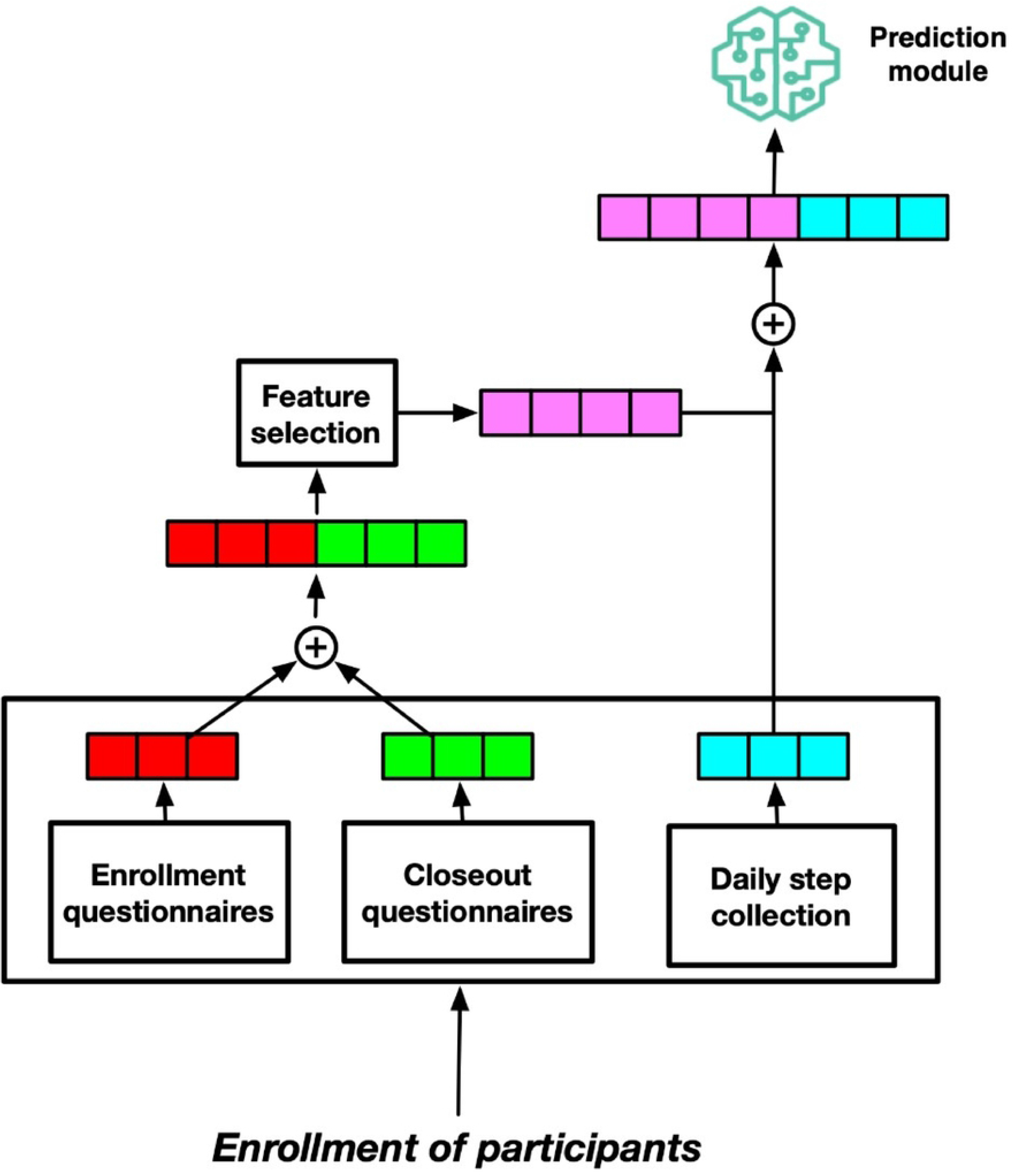
A schematic depicting study flow. At first, we combined the enrollment questionnaires and closeout questionnaires. Second, we performed feature selection step (including PCA, RFE, no feature selection, or features found from the previous study) with weekly steps data to develop a predictive model.

## Methods

### Data Collection

The data for this study was collected from a 9-week, non-randomized pilot study. For this purpose, 30 adults (9 male and 21 female) were enrolled from a local MGH-affiliated clinic with a body mass index (BMI) of 25 kg/m^2^ or greater. After screening the participants and seeking their consent, the research team directed the participants to the study website (http://www.wellocracy.com) to read through information regarding the study, the types and the features of available activity trackers during the 9-weeks study. The study staff assisted the participants with the device set up process as needed. In the study 27 participants chose to use the FitBit Charge model, two of them the FitBit One, and one of them chose the FitBit Zip.

Multiple surveys were collected from each participant at both enrollment and closeout stages: Behavioral Regulation in Exercise Questionnaire (BREQ-2) [16]; Barriers to Being Active (BBA) [17]); a depression screening Patient Health Questionnaire (PHQ-8) [18]); Prochaskas Stage of Change [19]; and general health questions (PROMIS Global-10) [20]).

These surveys included questions about their technology use and ownership, thoughts about and perceived barriers to exercise and activity (BREQ-2 and BBA), a depression screening (PHQ-8), Prochaskas Stage of Change, and PROMIS Global10. The BREQ-2 questionnaire is designed to gauge the extent to which reasons for exercise are internalized and self-determined based on the following categories: motivation, external, introjected, identified and intrinsic. In contrast, the BBA is designed to assess whether participants gauge certain categories as reasons for inactivity and include energy, willpower, time, resources, etc.

For each category, a score of 5 or greater would indicate that category as a substantial barrier to a person’s ability to exercise. During the study 10 participants were removed from the study since they did not use the trackers. That left the study with 20 participants.

Participants were instructed to continuously wear the activity tracker for the entire period of the study. The first week was treated as a run-in period. For the remaining weeks, the step goal for each participant was determined as 110% of their respective average step count achieved in the first week. Participants were contacted minimally during the eight-week period to facilitate observation of participants’ activity-tracker habits without interference.

At the end of the study, participants completed a closeout survey either online or in paper format and underwent a phone interview to gather information on their experiences during the study. All interviews were conducted and transcribed for analysis by a trained neuropsychologist.

### Experimental Setting

We divided the data into training and test sets. These sets are disjoint with respect to the participating patients. We chose 16 participants’ data as training dataset and remaining 4 participants’ data as testing dataset. We performed all the experiments and tuned hyper-parameters over training dataset. We defined the weekly target value as the average of steps per week for each participant. We did not consider weeks during which participants used the tracker device for less than four days.

### Data pre-processing

We collected all data from earlier mention questionnaires and the participants’ daily step count data from the activity trackers. These questionnaires generate 96 variables. The variables are screened to determine candidate predictors for building machine learning models. In the first iteration of the variable screening process, 11 variables, whose variance was zero, were eliminated. In the second iteration, we examined pairwise correlation among variables. We eliminated redundant variables in case Pearson correlation was above 0.9 for a variable pair. However, no pairwise correlation was found high enough to eliminate variables.

### Machine learning techniques

Figure 2 presents the techniques we have used to predict participants’ activity target point for the upcoming week. The selection of these methods is based on their strengths and capabilities that are not often overlapped. Bayesian ridge regression (BRIDGE) estimates a probabilistic model of a ridge regression [21]. LASSO is regression model that is robust to overfitting due to its regularization penalty [22]. RF is chiefly used for building predictive models, for both classification and regression problems. The RF can be interpreted easily and are less susceptible to underfitting [23]. NNs are modeled to mimic human brain. A NN consists of thousands or millions of neurons that perform mathematical operations designed to recognize patterns [24]. SVR is a regression model with advantage that its optimization does not depends on dimensionality of the data [25].

**Figure 2:**
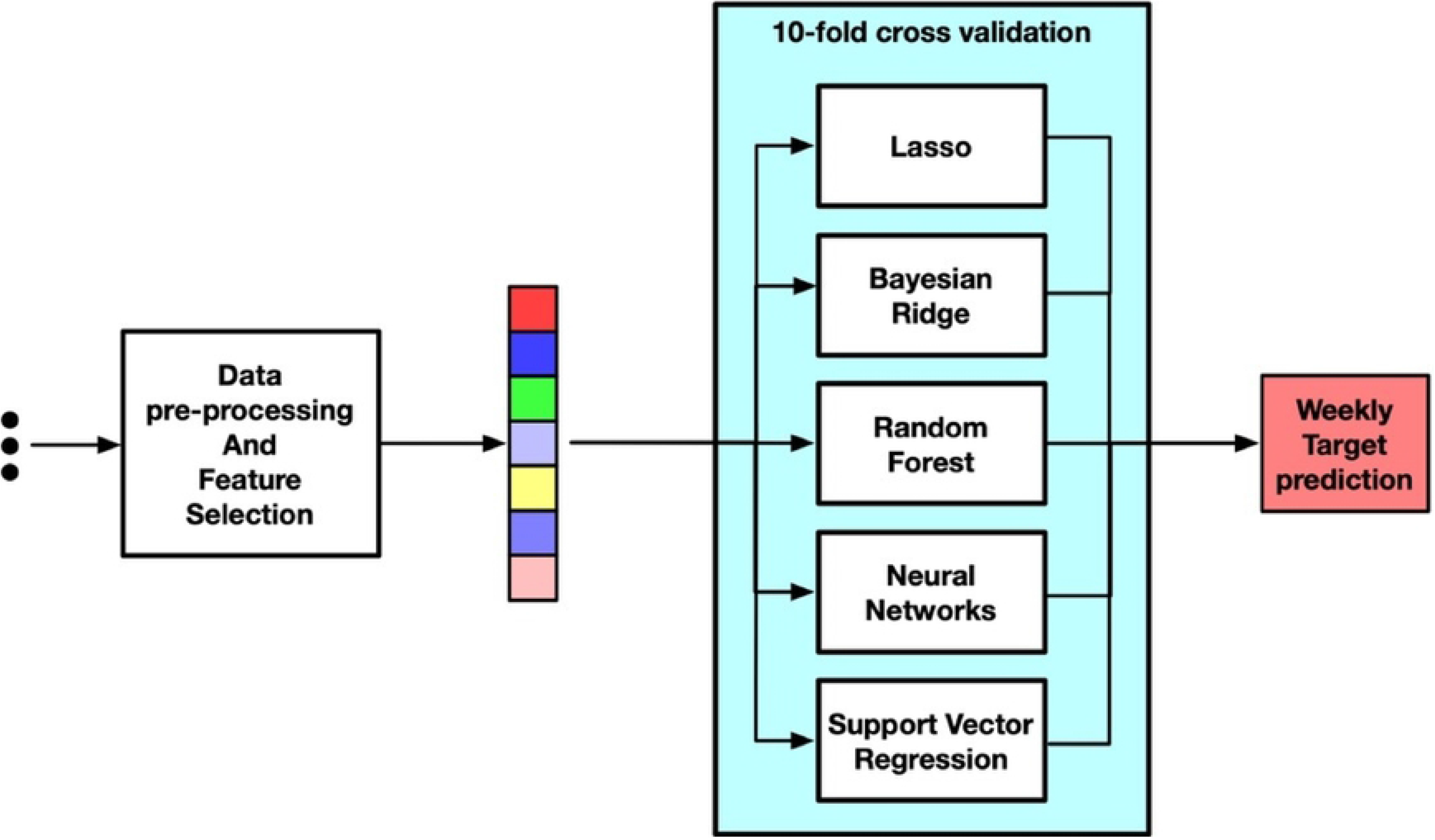
A schematic depicting models developed in the training step. We employed these machine learning models over feature extracted from (I) all features, (II) RFE features, (III) PCA features and (IV) features found in previous study.

We employed these models in four steps: (I) using all features, (II) using important features found by RFE (III) using new features developed by PCA and (IV) using subset of feature from authors’ previous study.

### Feature selection and extraction

In this study we were interested in using participants’ mental, behavioral information and weekly activity performance to predict the target step goal for the upcoming week. The questionnaires provide a highly redundant and low variance dataset. These features coupled with the daily step count which are aggregated on a weekly basis [Figure 3] and a normalized Shannon entropy (1) value of the weekly performance were considered candidate predictors of target step count for participants for the forthcoming week [26].

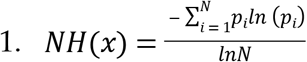

Where *i* denotes the day, *N* denotes the number of days per week (i.e., *N*=7) and *NH*(*x*) denotes the normalized Shannon entropy of the *N* daily step counts monitored during *N* days of a week. The normalized Shannon entropy varies between 0 and 1. A value close to 0 indicates that the step counts of the participant through the week are irregular; in contrast, a value close 1 indicates that the step counts are consistent over the week. In total there are 93 (=85+7+1) candidate predictors for building the predictive models. We used all 93 features for models developed in step I.

**Figure 3:**
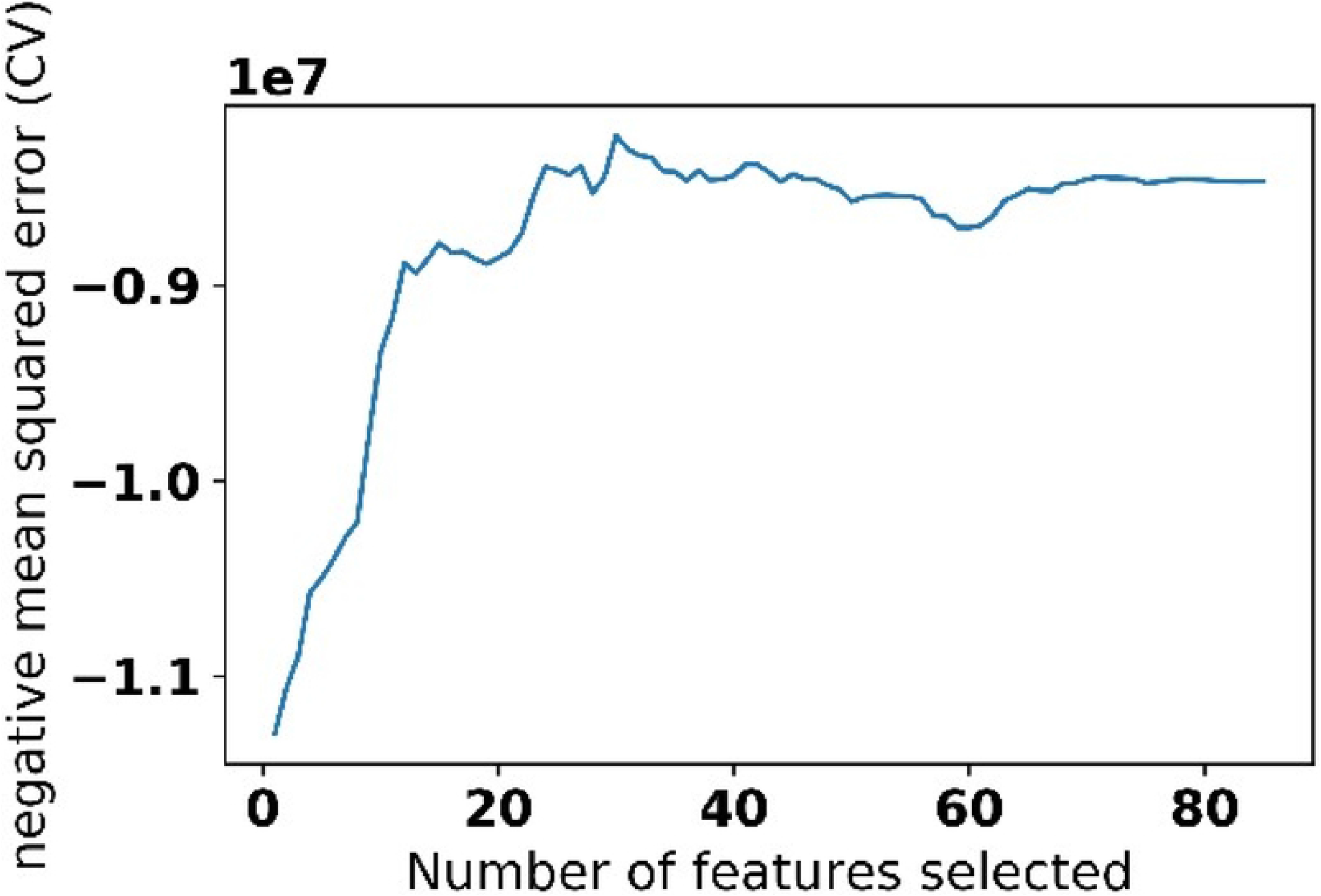
Daily steps distribution over 7 days in a week

In step II we performed a recursive feature elimination (RFE) with support vector regression over the 85 questionnaires’ variables to identify the important features. We augmented the important features from RFE model with the daily step count which are aggregated on a weekly basis and a normalized Shannon entropy value of the weekly performance were considered candidate predictors of target step count for participants for the forthcoming week. For step III, we used Principal component analysis (PCA) for dimension reduction. PCA is a dimensionally reduction technique that is widely used for extracting uncorrelated features components from correlated features.

We used PCA over the 85 features from the questionnaires. We combined the generated components with daily step counts and a Shannon entropy value of the steps counts. Lastly, for the step IV, we picked two variables from the questionnaires’ which were found to be important in a previous study. We coupled these two features with daily step counts and a Shannon entropy value of the steps counts.

### Model Evaluation

We trained the models developed in this study in a 10-fold cross validation using the training dataset. We compared the developed models using a MAE and adjusted-r^2^. We tested the final models of each step (**A, B, C and D models**) over an unseen testing dataset.

### Statistical Analysis

We used Python and R version 3.4.1 for model development and statistical tests. We performed a one-sided paired t-statistic (*α*=0.05) for statistical comparison between the models. The null hypothesis of the test was that the mean error of two models are equal with alternative hypothesis that a model (**best model**) has a lower mean error than the mean error given by a model under comparison with best model.

## Results

### Study cohort characteristics

We presented in Table 1 the characteristic of 30 participants of the study. Among all participants, 10 participants were removed from the study since they stopped using the activity trackers.

**Table 1.**
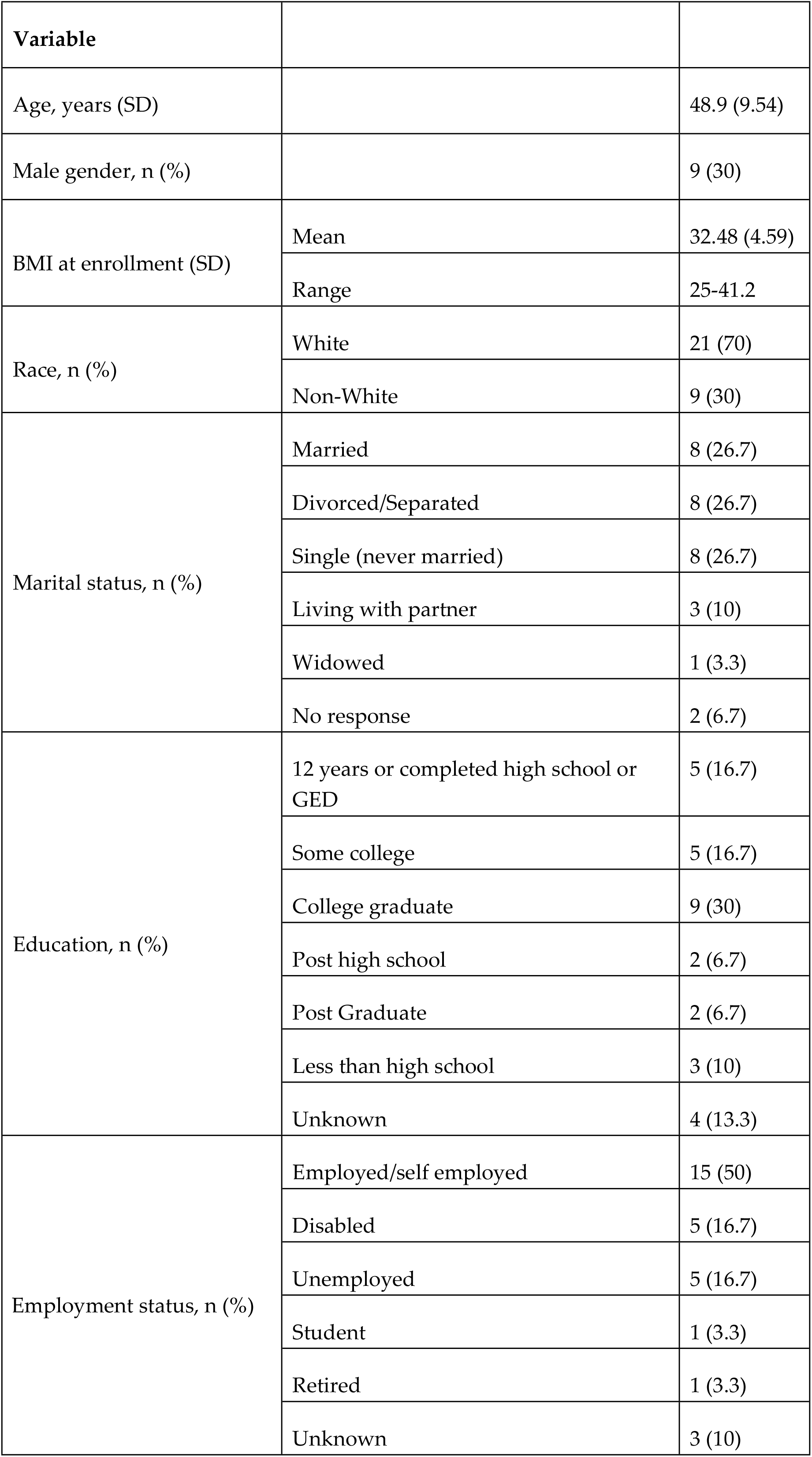
patient Demographic

| Variable |  |  |
| --- | --- | --- |
| Age, years (SD) |  | 48.9 (9.54) |
| Male gender, n (%) |  | 9 (30) |
| BMI at enrollment (SD) | Mean | 32.48 (4.59) |
|  | Range | 25-41.2 |
| Race, n (%) | White | 21 (70) |
|  | Non-White | 9 (30) |
| Marital status, n (%) | Married | 8 (26.7) |
|  | Divorced/Separated | 8 (26.7) |
|  | Single (never married) | 8 (26.7) |
|  | Living with partner | 3 (10) |
|  | Widowed | 1 (3.3) |
|  | No response | 2 (6.7) |
| Education, n (%) | 12 years or completed high school or GED | 5 (16.7) |
|  | Some college | 5 (16.7) |
|  | College graduate | 9 (30) |
|  | Post high school | 2 (6.7) |
|  | Post Graduate | 2 (6.7) |
|  | Less than high school | 3 (10) |
|  | Unknown | 4 (13.3) |
| Employment status, n (%) | Employed/self employed | 15 (50) |
|  | Disabled | 5 (16.7) |
|  | Unemployed | 5 (16.7) |
|  | Student | 1 (3.3) |
|  | Retired | 1 (3.3) |
|  | Unknown | 3 (10) |

### Parameter tuning and feature selection

We performed a grid-search with 10-fold cross validation over the training dataset. LASSO with alpha 0.01, BRIDGE with alpha 0.0001, SVR with polynomial kernel of degree 2 with C value of 0.00001 are the models considered in this study. For RF and NNs however the optimal models were different depending on the feature selection techniques. We found 30 variables from the questionnaires to be important using RFE with SVR [Figure 4]. Similarly, for PCA we found 14 as the optimal number of components that explains 100% of variance of the questionnaires as shown in Figure 5.

**Figure 4:**
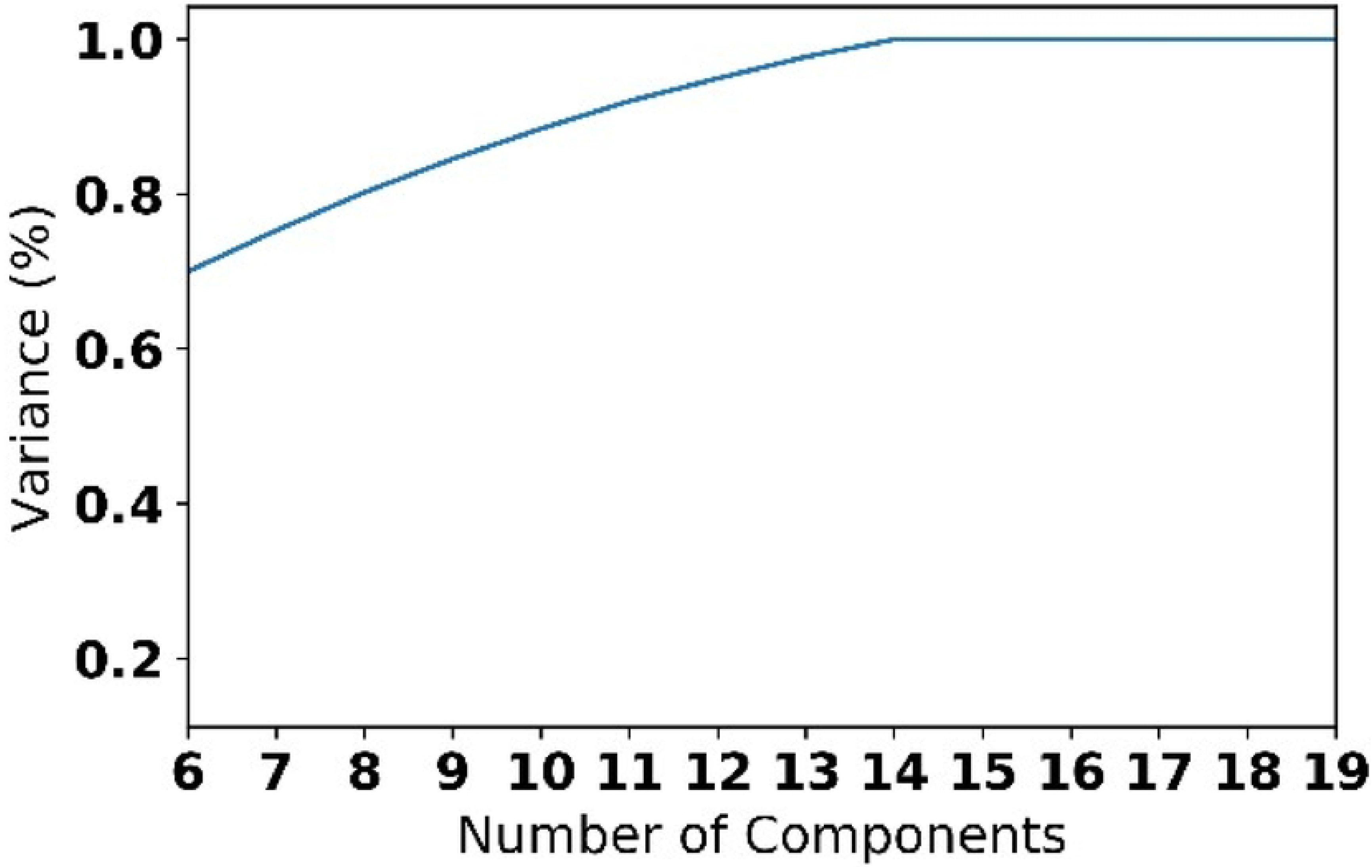
Number of features found to be important by SVR.

**Figure 5:**
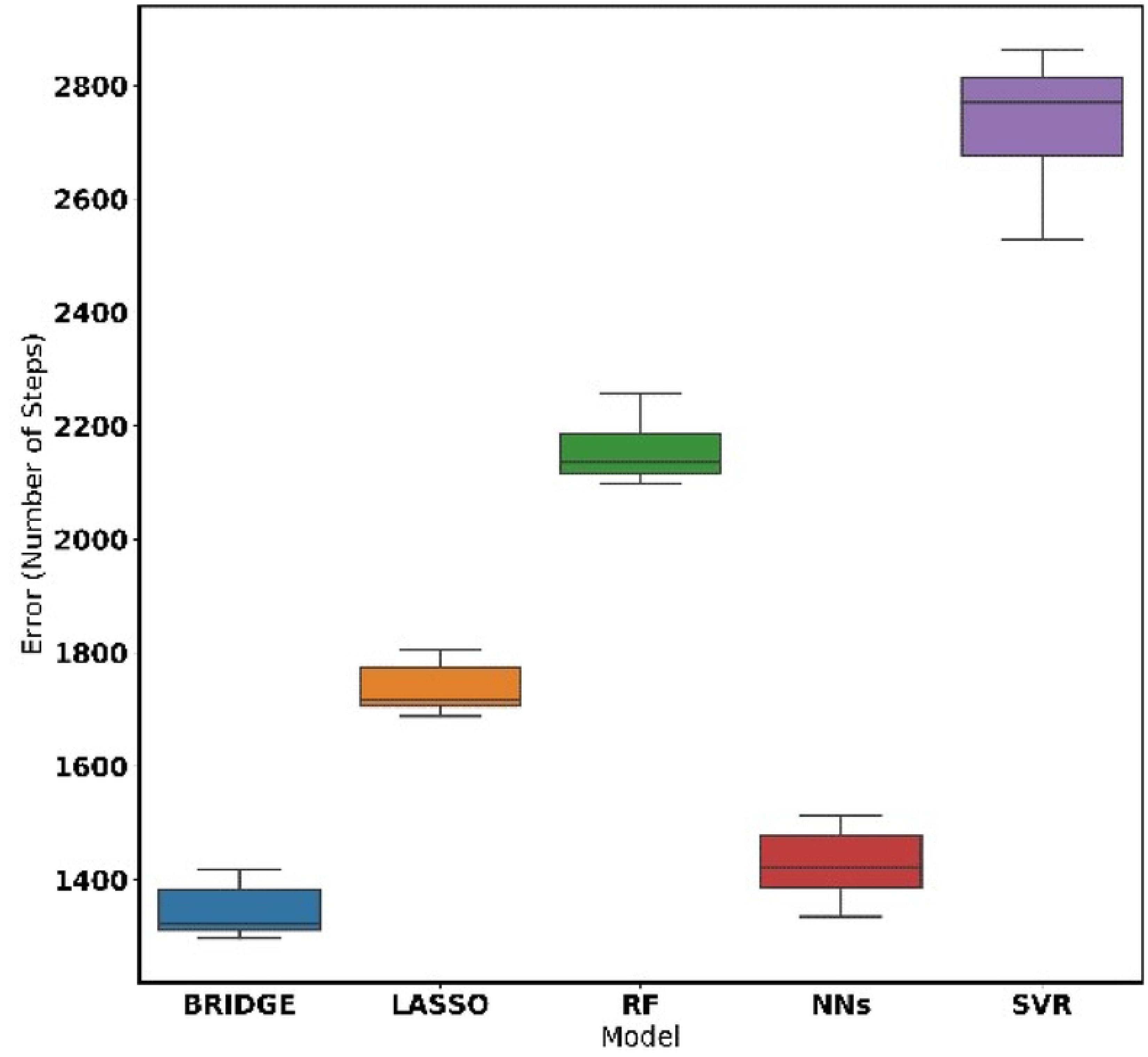
Number of components found by PCA.

#### Model performance

We reported the prediction performance of 10-fold cross validation for all models over training dataset in Table 2. We used mean absolute error (MAE) and *adjusted r*^2^ for the model comparison. Among all models developed, the BRIDGE model that uses 93 (=85+7+1) features (step I) gave the best performance over the training set [Figure 6]. It gave MAE of 1672 steps and *adjusted r*^2^ of 0.85. The BRDIGE model is referred as Model **A** in the rest of this paper. In step II, we coupled variables found by RFE with participants’ weekly performance features which resulted in 39 (= 30+7+1) features. A RF model with MAE of 1774 and *adjusted–r*^2^ of 0.81 had the best performance among all the models developed over these features [Figure 7]. We referred the RF model from this step as Model **B**.

**Figure 6:**
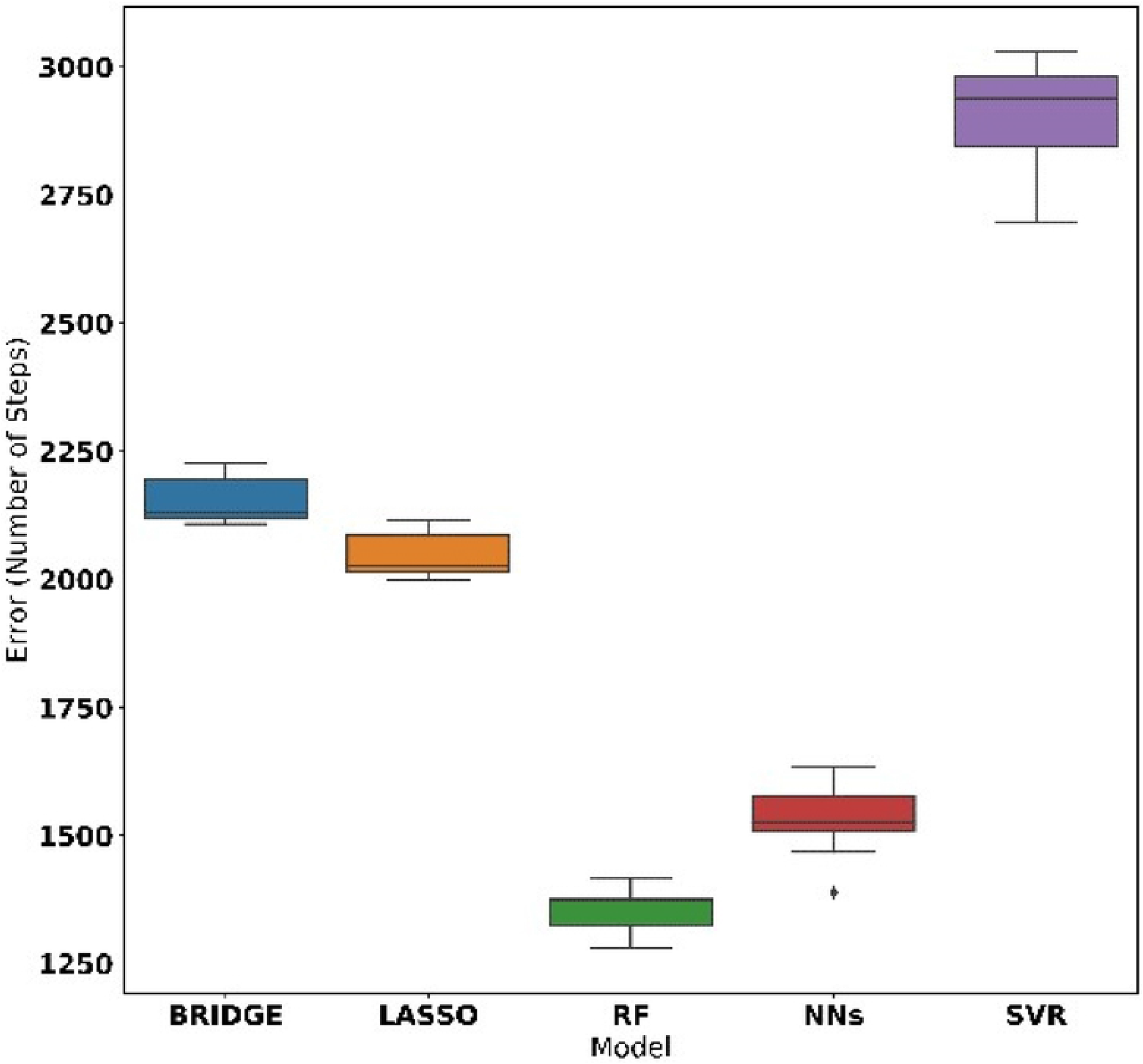
Performance models using all features plus daily step count of the current week and entropy of the daily of step count in the week

**Figure 7:**
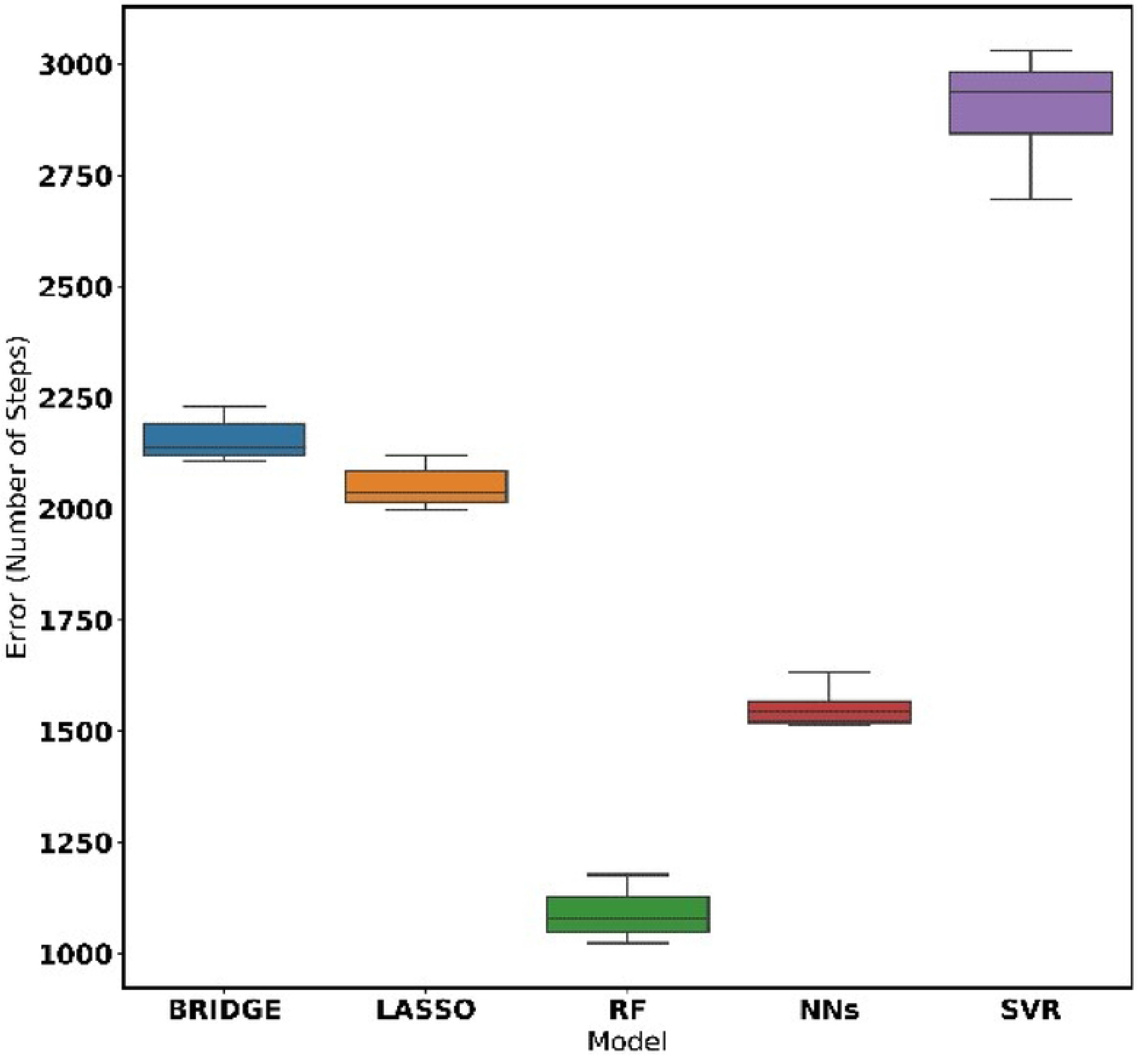
Performance of models using all features given by RFE plus daily step count of the current week and entropy of the daily of step count in the week

**Table 2.**
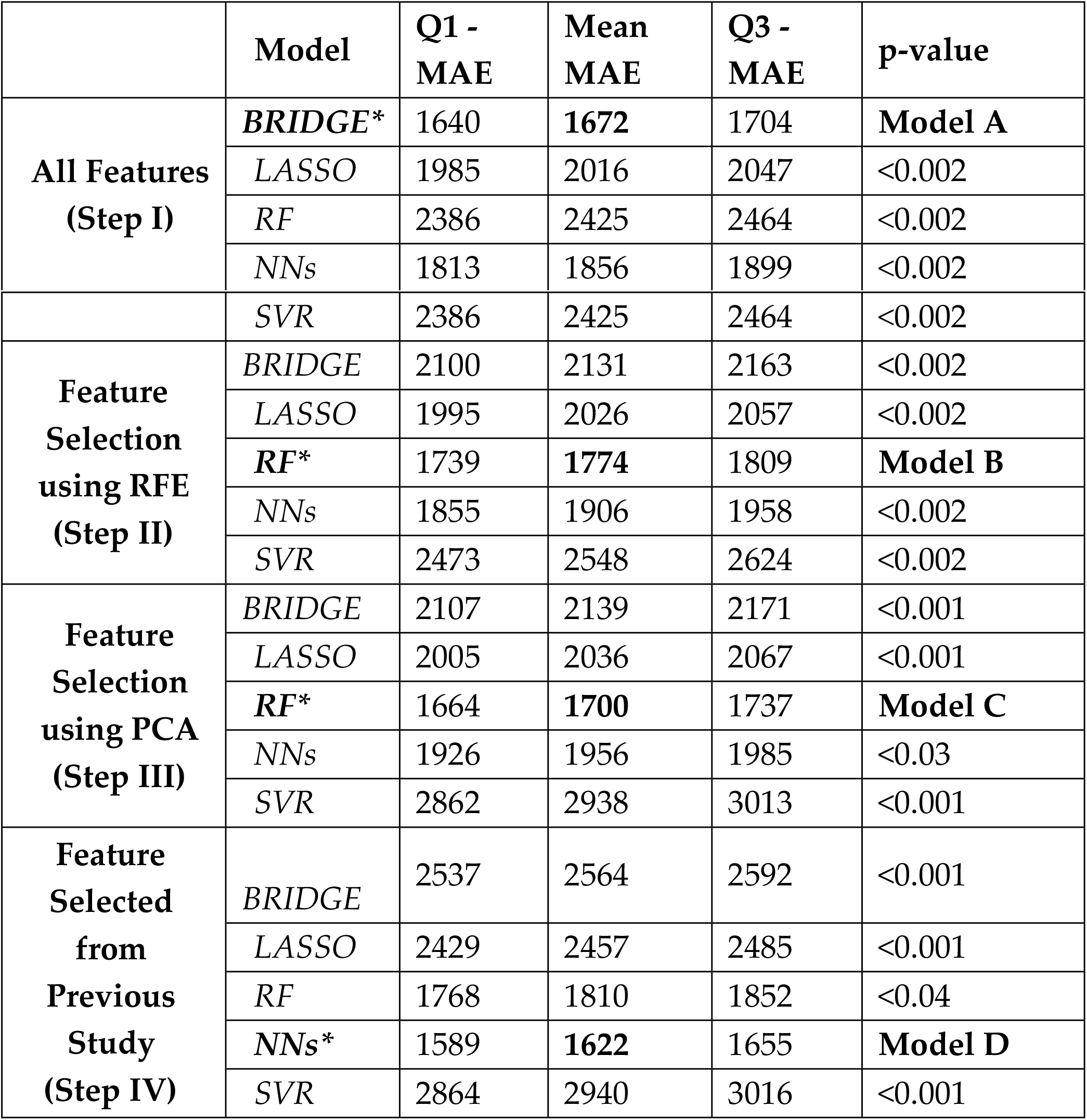
10 fold cross-validation results over training set,

Similarly, in step III, we coupled components extracted from PCA with participants’ weekly performance which resulted in 22 (=14+7+1) features. A RF with MAE of 1700 and *adjusted r*^2^ of 0.91 had the best performance among all models developed over these features [Figure 8]. This RF model is referred as Model **C** for the rest of this paper. Finally, for step IV, we coupled features found in the previous study with participants’ weekly performance which resulted in 10 (= 2 + 7 +1) features. NNs with the MAE of 1545 and *adjusted–r*^2^ of 0.77 gave the best performance across developed models over these features [Figure 9]. We referred to this NNs model as **model D**.

**Figure 8:**
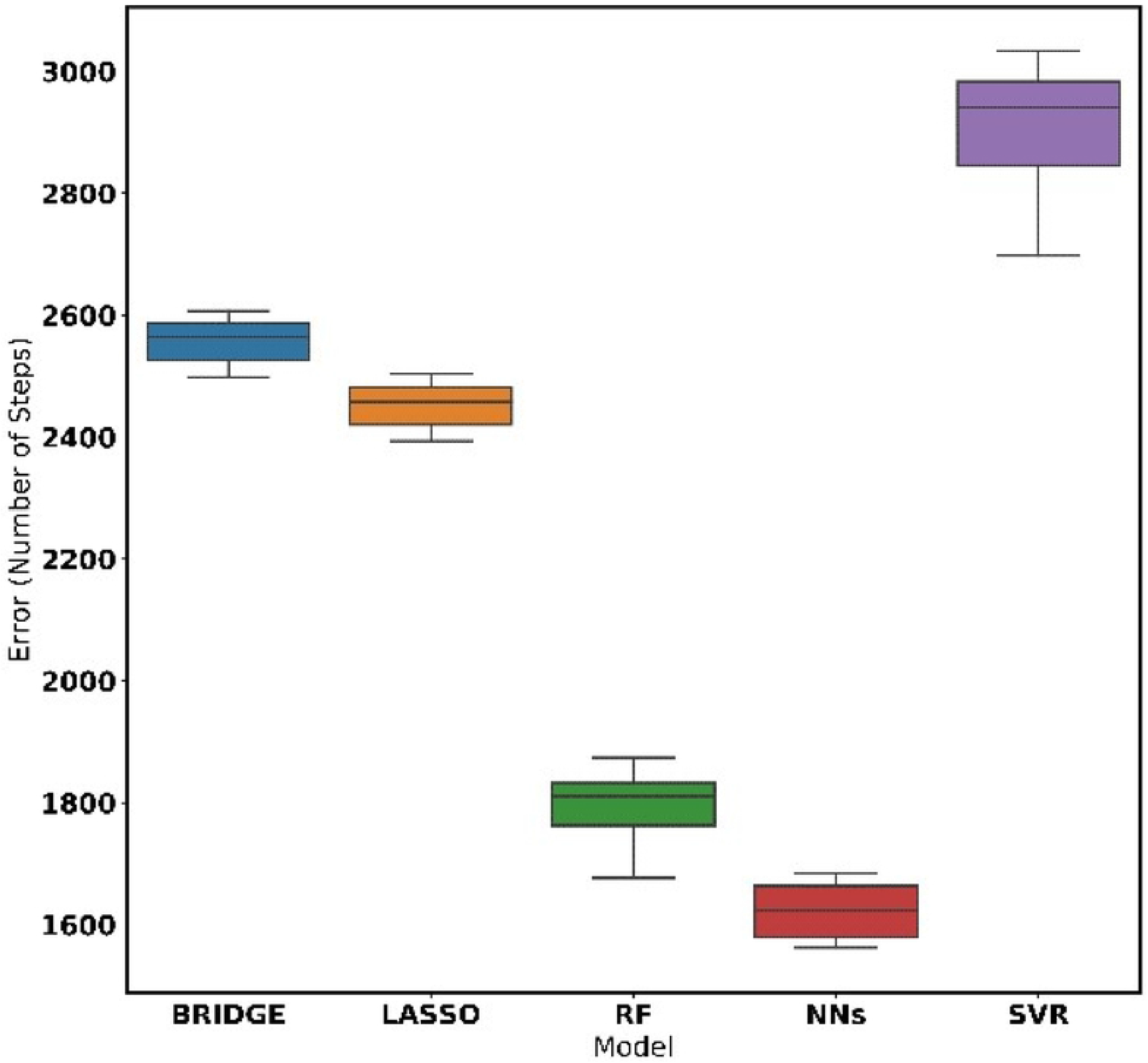
Performance of models using features generated by PCA plus daily step count of the current week and entropy of the daily of step count in the week

**Figure 9:**
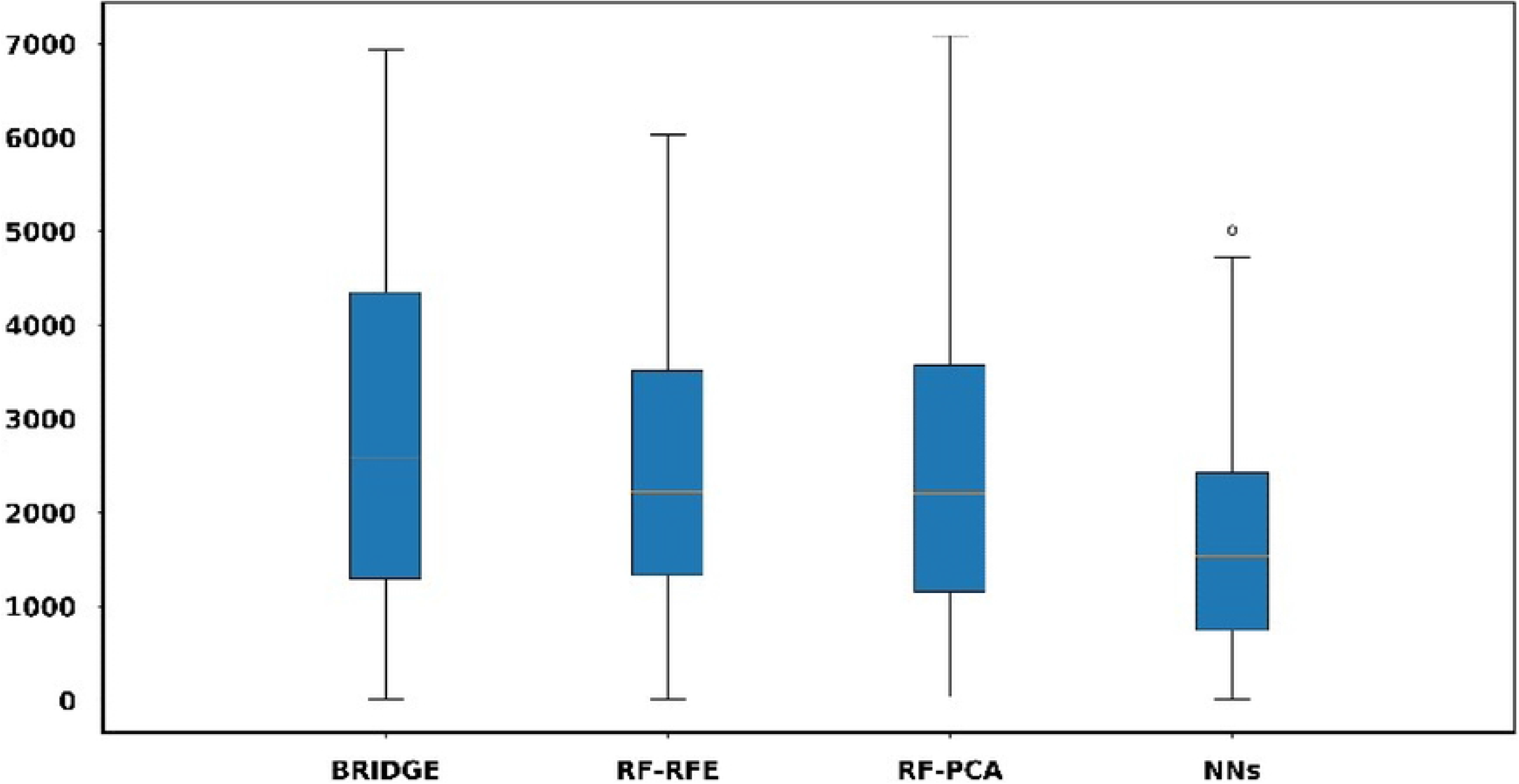
Performance of models using feature generated from previous knowledge plus daily step count of the current week and entropy of the daily of step count in the week

Model D had the lowest MAE across all developed models. We compared the predictive power of the **model D** with the best models [**A, B**, and **C**] from each step over the testing dataset (We found that the **model D** gives a better predictive performance for the testing dataset. We performed a t-test between the errors generated by the **model D** and errors generated by other models [**A, B, C**] as shown in Figure 10. We observed that the NNs (**D**) model has a lower error in comparison to the BRIDGE (Model **A** with p-value < 0.01), RF (Model **B** with p-value < 0.001) and RF (Model **C** with p-value < 0.01) as shown in Table 3.

**Figure 10:**
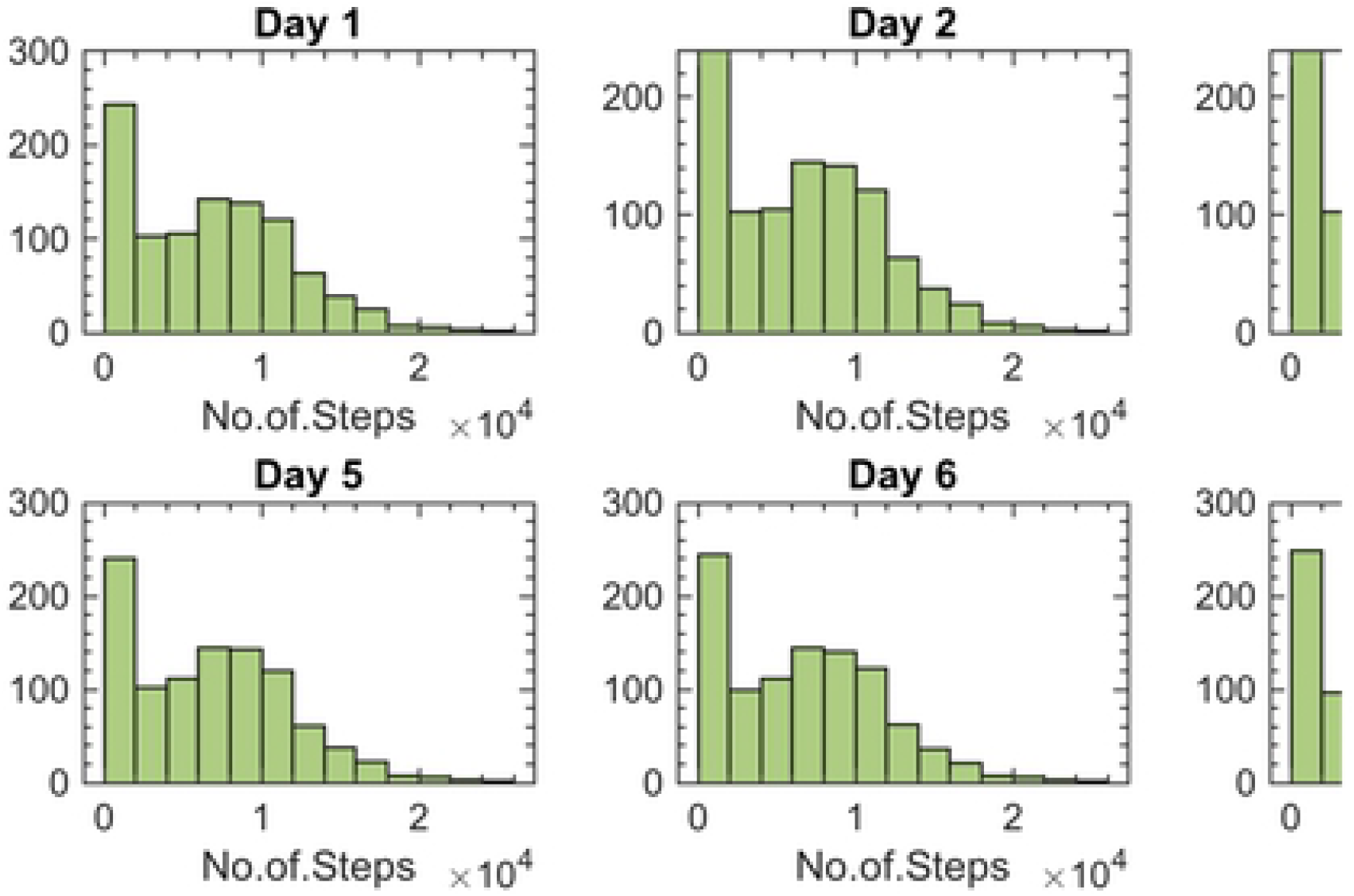
Performance of models - Best over test set

**Table 3.** Reference model results over test-set

| Model | Q1-MAE | Mean-MAE | Q3-MAE | P-value |
| --- | --- | --- | --- | --- |
| NNs ( model D) | 1383 | 1545 | 1706 |  |
| RF (Model C) | 1990 | 2210 | 2420 | <0.01 |
| RF (Model B) | 2015 | 2230 | 2445 | <0.01 |
| BRIDGE (Model A) | 2310 | 2578 | 2845 | <0.01 |

The **model D** achieved MAE of 1545 steps for 4 participants over a period of 8 weeks. This is a promising result considering that activity trackers are not accurate in daily activities [11].

## Discussion

To the best of our knowledge this is one of the first studies to develop a model to dynamically adjusted target number of steps based on an individual’s personal, social, environmental factors and weekly performance. Recent study concludes that activity tracker users feel unmotivated despite having an awareness of benefits of physical activity [27]. Users can become demotivated if they cannot meet their activity goals [28]. Easy-to-achieve goals may lead to abandonment of the activity tracker and setting too high step counts goals may discourage the individual from engaging in physical activity [29].

We studied the factors influencing the use of activity trackers and found that two factors promote the continued use of activity trackers: (1) the number of digital devices owned by the participant, and (2) whether or not the members of the participants’ family use activity fitness trackers and other smart devices. Extending from the previous work, in the current study, we developed a predictive model to estimate the target step goal for the upcoming week to encourage the use of activity trackers. The current study can be used the set goals for individuals accompanied by proper motivation to improve the sustained use of the activity trackers [30, 31, 32].

This study has some limitations. The number of participants was low, and all the participants were drawn from the same geographical area. Finally, participants who choose to participate in this study are more enthused about using Fitbit activity trackers than those who chose not to participate in the study.

To extend this work, a new study with the goal of validating the developed model with a large number of participants has been undertaken. The new study recruited 120 individuals from the general public to use the developed model over a period of 6 months. This model, which is hosted on a server, predicts a weekly target daily steps for each participant and updates the model parameters on a weekly basis with regard to the participants’ performance over the week.

## Code Availability

The code used for this analysis is available upon request, but it is not publicly available. Requests to access the codes will be reviewed by the project PI to verify whether restrictions due to Intellectual Property apply.

## Data Availability

The data that supports the findings of this study will not be publicly available due to confidentiality of the data.

## Acknowledgment

This research was supported in conjunction with a grant from the Robert Wood Johnson Foundation (Grant ID: 71963).

## Author Contribution

Mohammadi.R and Kamarthi.S worked on data exploration, dimension reduction through factor analysis, and implementing and designing neural network-based algorithm and generating results and preparing the draft of the paper. He is one of the guarantor authors of the paper.

